# CRP-Mediated Coherent Type-4 Feed-Forward Loop Involving Glucose-Regulated LacI in *Escherichia coli*

**DOI:** 10.1101/2025.11.14.688166

**Authors:** Yang-Chi-Dung Lin

## Abstract

Recent studies have highlighted the importance of structure-function relationships in genetic regulatory networks, particularly in feed-forward loops (FFLs), where input-output behavior depends on both input signals and transcriptional interactions. This study elucidates the function of a CRP-LacI-*lacZYA* coherent type-4 FFL (Co-4 FFL) in *Escherichia coli*. We demonstrate that cyclic AMP receptor protein (CRP) directly represses the transcription of *lacI*, which encodes the Lac repressor. This finding was confirmed through multiple approaches: 1) the mRNA level of *lacI* decreased to one-fifth of the original level upon cAMP addition; 2) *lacI* expression increased 15-fold in a *crp* mutant; 3) DNase I footprinting identified a CRP binding site within the *lacI* promoter region, and a lacZ fusion assay and site-directed mutagenesis validated its functional role. Collectively, these results establish CRP as a direct repressor of *lacI*, thereby forming a Co-4 FFL with the *lacZYA* operon. Physiological studies on Co-4 FFLs in *E. coli* are scarce. Our model suggests that under glucose-lactose diauxic growth conditions, this circuit allows *E. coli* to enhance the efficiency of lactose metabolism for energy synthesis upon glucose exhaustion by repressing *lacI* via CRP. This work reveals how metabolic cues shape the behavior of genetic regulatory networks.

## Introduction

Recent studies have yielded insights into structure–function relations in genetic regulatory networks. Models of feed-forward loops (FFLs) show that the input–output behavior depends critically on both the input signal and transcription interactions. Engineered systems are often composed of recurring circuit modules that perform key functions. Transcription networks that regulate the responses of living cells were recently found to obey similar principles. They contain several biochemical wiring patterns, termed network motifs, which recur throughout the network. One of these motifs is the feed-forward loop (FFL). The FFL, a three-gene pattern, is composed of two input transcription factors, one of which regulates the other, both jointly regulating a target gene. The FFL has eight possible structural types, because each of the three interactions in the FFL can be activating or repressing. In *E. coli,* coherent FFLs function as a sign-sensitive delay element in an *ara* system [1], and incoherent FFLs accelerate the response time of a *gal* system [2].

Conducting a large-scale microarray experiment, we found that different concentrations of cAMP (0 mM to 10 mM) added to the medium enhanced the effects of the cAMP-CRP complex (*Unpublished experimental data*); as a result, the gene expression of the LacI repressor (lacI) in the *Lac* operon was significantly inhibited. The cAMP-CRP complex, formed by cAMP and cAMP receptor protein (CRP) in E. coli, can function as a transcription factor, thereby affecting gene regulation and expression. Here, the function of a *CRP-LacI-lacZYA* coherent type 4 feed-forward loop (Co4-FFL) in *E. coli* was demonstrated by determining the regulatory role of CRP on LacI repressor and the Lac Operon.

The *lac* operon, also known as the lactose operon, is an operon essential for the transport and metabolism of lactose in *E. coli* and other enteric bacteria. It has three adjacent structural genes, *lacZ*, *lacY*, and *lacA*. The genes encode β-galactosidase, lactose permease, and thiogalactoside transacetylase, respectively [3]. In its natural environment, the *lac* operon allows for the effective digestion of lactose. The *lac* operon is a classic model of gene regulation in molecular biology, developed from *E. coli*. This control module enables *E. coli* to optimize the metabolism of carbohydrates, such as lactose, from its environment. For example, *E. coli* utilizes glucose as its primary carbon source in energy metabolism within environments rich in glucose. However, a large amount of lactose is present in the same environment [4–6].

In response to environmental changes, bacteria have evolved sophisticated regulatory networks, including multiple transcription factors (TFs), to adjust gene regulation in response to various environmental signals, such as the availability of carbon sources. For example, if glucose and lactose were both provided, glucose was metabolized first (growth phase I) and then lactose (growth phase II). Lactose was not metabolized during the first part of the diauxic growth curve because β-galactosidase was not made when both glucose and lactose were present in the medium. Monod named this phenomenon diauxie [7]. Until now, it is well known that the bi-phasic growth curve is correlated with intracellular cAMP levels and regulated by two transcription factors, CRP and LacI.

Diauxic growth occurs when microorganisms are in an environment with two different carbon sources [8, 9]. These microorganisms initially decompose the acceptor that can be easily consumed as an energy source, thereby inhibiting the consumption of the other acceptor. This phenomenon is called catabolic repression. After the easily used acceptor is completely decomposed, an intermittent period of growth occurs. During this period, a thallus induces a required enzyme by using the other acceptor to start another cycle of growth [10]. Studies have been conducted using glucose and lactose as carbon sources. *E. coli* uses glucose as the main source of energy. At this moment, lactase and related genes are inhibited. Therefore, this phenomenon is called glucose catabolic repression [11]. Further studies have investigated the mechanism by which cells induce lactase generation, revealing that glucose catabolic repression is closely related to the concentration of cAMP in cells [12]. As glucose is consumed gradually, the concentration of cAMP in cells likely increases gradually. cAMP and CRP form the cAMP-CRP complex, and this complex is combined with the operator area of the lac operon [13]. cAMP-CRP complex can also increase the affinity of RNA polymerase to DNA, thereby causing the overexpression of enzymes involved in lactose metabolism. On the contrary, low concentrations of cAMP form only an insufficient cAMP-CRP complex in glucose-rich environments; thus, RNA polymerase binding sites on the *lacZ* gene promoter of *the lac* operon are occupied by LacI. As a result, the expressions of a series of downstream lactase genes are inhibited.

In this study, an investigation was conducted to elucidate the regulatory mechanism by which glucose affects the expression of the *lacI* gene and *lacZ* gene. Our results showed that the expression of the *lacI* gene changed significantly when *E. coli* was cultured in LB with glucose or with cAMP. After glucose was added, the mRNA expression of *lacI* increased by a factor of three. Conversely, the mRNA expression of *lacI* decreased to one-fifth of the original value after cAMP was added. Based on these experimental results, we found that this phenomenon may be controlled by the transcription factor CRP. Therefore, we initially investigated the activity of the cAMP-CRP complex on the transcriptional expression of the *lacI* gene and further evaluated the physiological mechanism resulting from this regulatory effect. We also aimed to integrate the obtained results and establish the regulatory relationship between cAMP-CRP and *lacI* in relation to the regulation of the *lacZ* gene and *lac* operon.

We express and purify the CRP protein and conduct an in vitro assay test (EMSA) using this promoter. DNase I footprinting was performed to analyze the known binding area of CRP and identify the binding sequence of CRP on the *lacI* promoter. We also repeatedly conducted this experiment using a CRP mutant and a *cyaA* mutant strain to verify whether the effect on lacI expression is related to CRP being affected by glucose. Furthermore, we used an antibody against the LacI protein in a Western blot to detect the expression of the LacI protein under the previously described experimental conditions. These procedures were conducted to investigate whether the glucose-induced change in lacI gene expression can be determined at the transcriptional level.

After re-establishing the regulatory relationship between CRP and LacI with respect to the *lac* operon, we determined the function of CRP-LacI-*lacZYA* Co-4 FFL. Above all, we suggest that *E. coli* uses CRP to repress *lacI* expression, thereby increasing the efficiency of metabolizing lactose for energy synthesis under glucose exhaustion in glucose and lactose diauxic growth conditions. The hypothesis was further verified by site-directed mutation of the CRP binding site on the *lacI* promoter region. We then analyzed whether wild-type and mutant strains can alter the expression level of *lacZ*. This experiment was also performed to verify whether or not CRP can control the application model of *E. coli lacZ* gene regulation in an environment containing double-carbon sources (glucose and lactose) with Co4 FFL formed by regulating the *lacI* gene.

## Experimental Procedures

### Bacterial strains

Mutant strains used in this study were *E. coli* BW25113 derivatives generated from the Keio collection system, which were provided by the National Institute of Genetics in Japan [14].

### Plasmid construction

The *lacI*::*lacZ* reporter plasmid, pLACI, was constructed by introducing the PCR product with the 500 bp region upstream to *lacI* start codon into pRW50, a low-copy-number *lacZ* expression vector (provided by Dr. Busby of the University of Birmingham) [15]. The pCAN plasmid carrying a mutant CRP binding site was derived from pLACI by site-directed mutagenesis to verify the function of the predicted CRP binding site on the *lacI* promoter region.

### Electrophoretic mobility shift assays

EMSAs were conducted as described in previous studies [16–18] with some modifications. DNA fragments for EMSA were obtained by annealing commercially synthesized complementary oligonucleotides (by heating to 65 °C and then cooling to room temperature) or by PCR. The EMSA reaction mixtures (20 µl) contained DNA oligonucleotides (20 nM), cAMP (200 μM), and different concentrations of CRP (0, 0.05, 0.1, 0.2, 0.4, and 0.8 µM) in a buffer solution of Tris-HCl (20 mM, pH 8.0), MgCl_2_ (10 mM), ethylenediaminetetraacetic acid (EDTA, 0.1 mM), and KCl (100 mM). The reaction mixtures were gently mixed and incubated for 30 minutes at 37°C to allow for binding. Novel juice loading dye (GeneDireX, Las Vegas, NV, USA) was subsequently added to the reaction mixtures and separated by non-denaturing polyacrylamide gel (6%). To confirm the predicted CRP binding site on the *lacI* promoter, a DNA fragment carrying the CRP binding sequence upstream of the *lacI* gene was synthesized to confirm the binding ability of the cAMP–CRP complex. The DNA fragment (NCA fragment) containing the mutated CRP binding site served as a negative control.

### DNase I footprinting

DNase I footprinting assays were performed as described previously [16, 19–21]. The PCR product carrying the –350 to –161 region of the *lacI* promoter was generated by 5’FAM-labeled primers. The binding of the cAMP–CRP complex to the labeled DNA fragment was conducted as described for EMSA. The binding mixtures were then partially digested with 0.4 units DNase I (Promega, Madison, WI, USA) in the absence or presence of CRP (400 nM). After incubation for 45 s at 37 °C, the reaction was stopped by heating at 98 °C for 10 min without the addition of EDTA. The digested DNA fragments were purified using the MinElute PCR purification kit (Qiagen, Hilden, Germany) and then separated by capillary electrophoresis using the Applied Biosystems 3730 DNA analyzer (Applied Biosystems, Foster City, CA, USA). The protective regions of cAMP–CRP DNA binding to DNase I digestion were analyzed using the Peak Scanner Software v1.0.

### Quantitative mRNA expression level

RNA manipulation and real-time PCR were performed as described previously [16, 22]. RNA isolation is based on the suggested protocol in the TRI reagent-RNA kit (Molecular Research Center, Cincinnati, OH, USA). RNA samples for real-time PCR were pre-treated with DNase I (Promega, Madison, WI, USA). The DNA primers (listed in Table S2) used in real-time PCR were designed using Primer Express software (Applied Biosystems, Foster City, CA, USA), and the complementary DNA was synthesized with SuperScript III Reverse Transcriptase (Invitrogen, Carlsbad, CA, USA). Real-time PCR was performed using the ABI PRISM 7000 (Applied Biosystems, Foster City, CA, USA) and SYBR Premix Ex Taq (Takara, Tokyo, Japan). All quantitative PCR reactions were performed with six replicates.

### Promoter activity assay

The activity of β-galactosidase was determined in *an E. coli* BW25113–derived *crp* null strain. The mutant strains were co-transformed with reporter plasmids (pLACI and pCAN) and pDCRP, which encodes wild CRP. The pLACI plasmid has a wild-type lacI promoter. In contrast, the pCAN plasmid contains a mutation at the CRP binding site. The cells were grown in LB at 37 °C until the optical density (OD) at 600 nm (OD_600_) reached 0.4. All assays were performed with five replicates.

### Site-directed mutagenesis of the CRP binding site

The promoter region of lacI was cloned into a TA vector (pGEM-3 Easy vector system, Promega) and transformed into DH5α competent cells. Point mutation primers were designed for mutagenesis by using QuikChange Primer Design (Agilent). The TA vector was PCR amplified using the point mutation primer. The PCR program was set as follows: 94℃ for 2 min was used to denature the DNA double helix. 15 cycles (98 ℃, 10 sec; 61℃, 30 sec; 68℃, 3min 50 sec) was used to amplify plasmid. And then, a final extension of 72℃ for 7 min was used. Finally, the temperature dropped to 4℃. DpnI digested the PCR product of the TA vector for 1 hour. The five μL final product was transformed into 50 μL DH5α competent cells by heat-shock transformation. The transformed DH5α competent cells were plated onto an LB plate containing ampicillin (100 μg/mL). Colonies on the plate were cultured, and the cultures were used to extract the plasmid. Sequencing of the plasmid was applied to check the mutagenesis result.

### Site-directed mutation on the *Escherichia coli* chromosome

Site-directed CRP binding site mutations on the *lacI* promoter region on the *E. coli* chromosome were conducted by Counter-Selection BAC Modification Kit [23]. Before using this kit, the BW25113 rpsL* streptomycin-resistant strain must be prepared first, by carrying the mutated rpsL gene from the HS996 strain into the BW25113 strain through P1 transduction. And then the BW25113 strain, carrying a mutated rpsL gene, was transformed with the pRed/ET plasmid, which encodes recombinase. Generation of *the rpsL-neo gene flanked by lacI homology regions* was done by PCR with the. This PCR product was transformed into cells carrying the pRed/ET plasmid, which was induced by arabinose to facilitate recombination. The successful colony could resist kanamycin. Next, the non-selectable DNA, lacI* promoter region, was amplified from TA_lacI * and transformed into the cells obtained in the previous step. The *rpsL_neo* cassette was replaced by a non-selectable DNA sequence, including the lacI promoter region, through recombination. It led the cells to lose resistance to kanamycin and recover resistance to streptomycin. The result of cloning was check by sequencing.

## Results

### *lacI* transcription is negatively regulated by the cAMP–CRP complex

CRP is a well-known transcription factor governing carbohydrate metabolism [24]. When *E. coli* grows in a medium containing glucose, intracellular cAMP concentrations are low [25]. To investigate whether the glucose effect on lac operon gene expression is associated with the CRP-LacI-mediated feed-forward loop, real-time PCR was performed to measure mRNA expression of lacI, which encodes the LacI repressor, in wild-type and *crp* deletion mutant cells. We found that the abundance of the *lacI* transcript increased threefold in the presence of glucose and was positively correlated with glucose concentrations (**Figure 1**). In the CRP deletion mutant cells, *lacI* expression was not affected by altering the glucose concentration in the medium. This result suggests that CRP is a mediator linking glucose concentration to lacI expression, and that this expression is not caused by the indirect effects of high glucose concentration in the medium.

**Figure 1.**
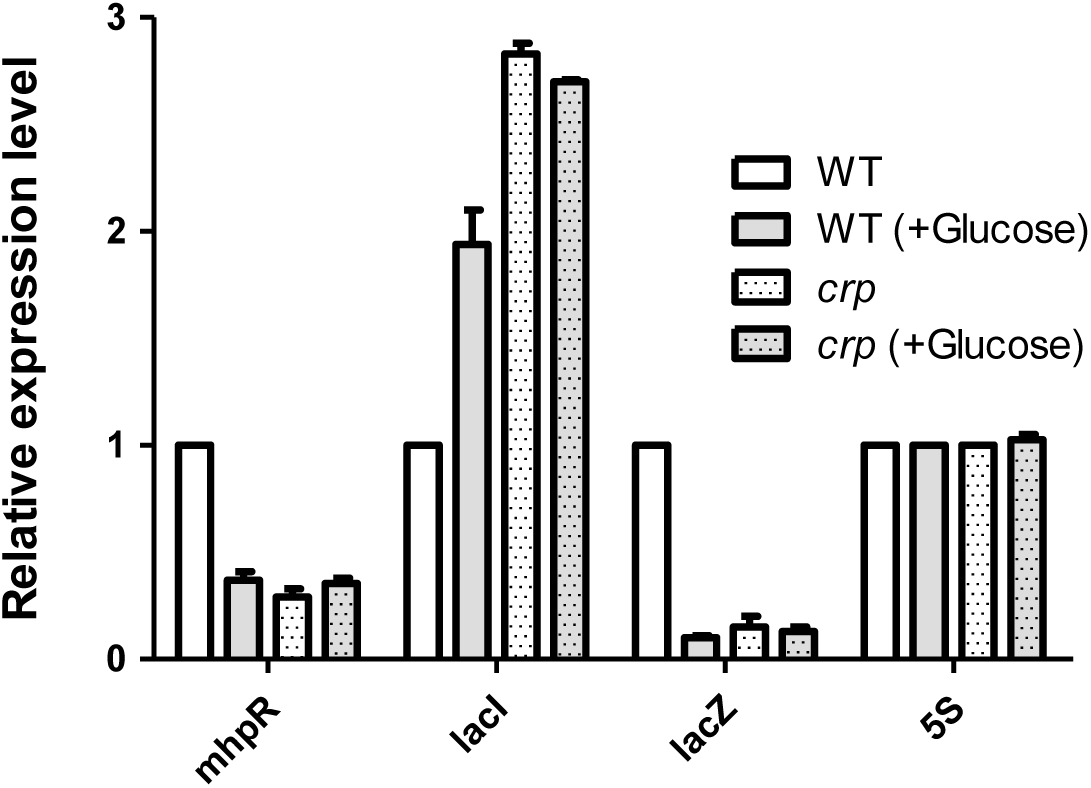
Effect of CRP and glucose on cAMP–CRP-dependent mhpR, *lacI* and *lacZ* genes regulation. BW25113 and its isogenic *crp strains were cultured in LB or LB with 40* mM glucose. In the exponential phase, cells were harvested for RNA extraction and further analysis. The RNA expression level was detected by qRT-PCR and normalized to that generated from LB-cultured wild-type (wt).

The DNA-binding ability of CRP is regulated by cAMP concentrations [24, 26]. Therefore, we determined whether CRP can regulate *lacI* operon expression under different cAMP concentrations (0 mM or three mM). A quantitative measurement of CRP-mediated transcriptional regulation was performed by measuring the *lacI* gene expression level in an *E. coli* K12 strain lacking the *cyaA* gene, which produces endogenous cAMP [27], under different cAMP concentrations. mRNA levels of the *lacI* gene decreased by 20% in LB medium with cAMP at three mM (**Figure 1**). In contrast, in the CRP deletion mutant cells, transcription of the *lacI* gene increased compared with the transcription level in the wild-type cells (**Figure 1**). The *mhpR* gene, known to be regulated by CRP and located upstream of the lacI gene, served as a control group in this experiment, thereby ensuring the validity of the experimental results. These results suggest that CRP acts as a repressor of the *lacI gene*.

The regulatory role of the cAMP-CRP complex on *lacI* gene expression was investigated under different nutritional conditions (**Figure 2**). In the wild-type strain (BW25113) grown in LB medium alone, a baseline expression level was established. The addition of 10 mM glucose to the LB medium led to a significant decrease in lacI mRNA levels. Conversely, supplementation with three mM cAMP resulted in a marked increase in lacI expression. To genetically validate this dependence, we analyzed Δ*crp* (lacking CRP) and Δ*cyaA* (lacking adenylate cyclase, thus unable to produce cAMP) mutants. In both mutant strains, *lacI* expression was severely impaired under all conditions tested (LB, LB with glucose, and LB with cAMP), remaining at a very low level comparable to the glucose-repressed state in the WT. These results demonstrate that the cAMP-CRP complex acts as an essential positive transcriptional regulator of the lacI gene.

**Figure 2.**
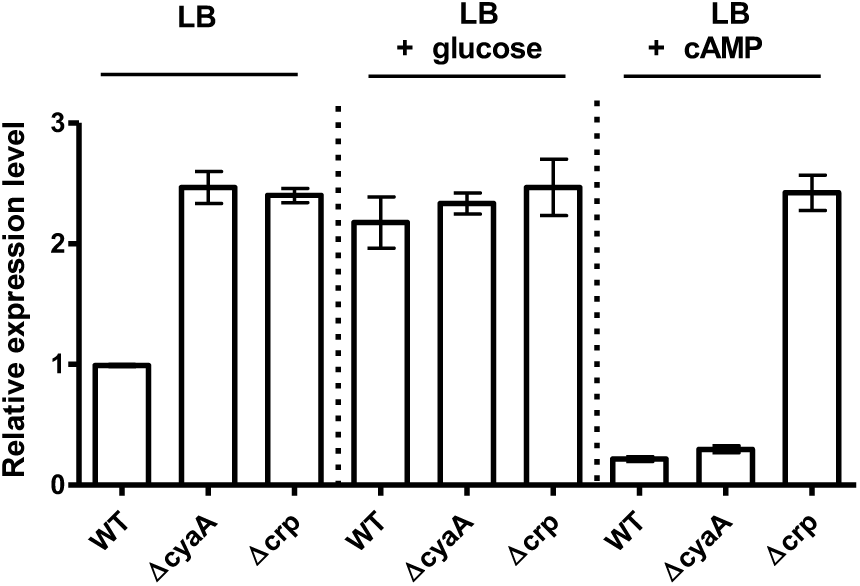
cAMP-CRP-dependent regulation of *lacI* gene expression. lacI expression was quantified by real-time PCR in WT, Δcrp, and ΔcyaA strains grown in LB with the indicated supplements. Data (mean ± SD) are normalized to 16S rRNA and expressed relative to the untreated WT control.

### Identification of the CRP-binding site in the *lacI* promoter region

To identify potential CRP-binding sites in the *lacI* promoter region, EMSA and footprinting analyses were performed [16, 19–21]. EMSA was performed first to test the interactions between the cAMP–CRP complex and three sequential fragments located from –350 to +100 bp relative to the *lacI* transcription start site (**Figure 3A**). The only fragment showing retarded migration by the cAMP–CRP complex was the fragment covering the –50 to +100 bp upstream region. To precisely locate the CRP-binding site, the *lacI* promoter region was scanned using the MATCH program [28] to predict the location of the cAMP–CRP complex binding site. Bioinformatics analysis revealed a potential CRP-binding site between −5 and +20 bp of the *lacI* promoter region. The predicted CRP-binding site was further verified by EMSA (**data not shown**). In a typical procedure, a *lacI* fragment carrying the −25 to +25 bp of the *lacI* gene was synthesized to confirm the binding ability of the cAMP–CRP complex. Retarded migrating bands became evident with increasing CRP concentrations, whereas the LacIP* fragment (change the consensus sequence of CRP binding site) exhibited no CRP-binding ability (**Figure 3B**). We also conducted DNase I footprinting to map the CRP-binding site in a *lacI* promoter fragment covering –100 to +100 bp. Comparison of the sequence patterns in the absence and presence of CRP (0.4 µM) revealed that only one protein-protected region covering the *lacI* promoter region from −6 to +22 bp was protected by CRP (**Figure 3C**). Protective regions of the *casA* promoter were approximately 30 bp long.

**Figure 3.**
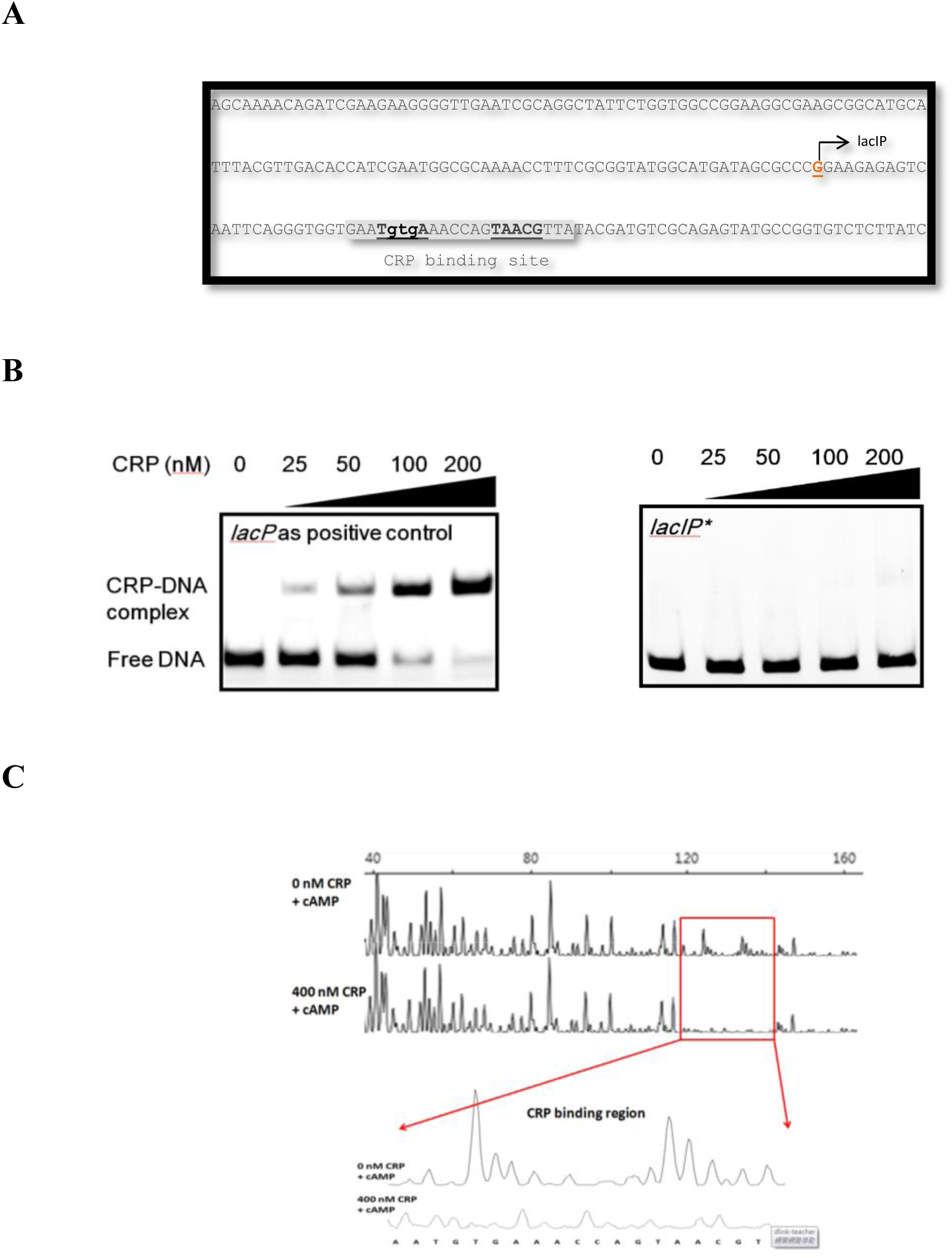
Identification of the CRP-binding site in the lacI promoter region. **A.** Sequence map of the LacI promoter region. The sequence of the *lacI* promoter region. The CRP site is indicated with a black-shaded gray area. **B.** The mutated CRP binding site on the lacI promoter region has no CRP binding ability. **C.** DNase I footprinting. Mapping the CRP binding site on *the lacI* promoter by DNase I footprinting.

### Construction of an *E. coli* Strain with a Mutated CRP-binding Site in the *lacI* **Promoter**

To investigate the specific role of the cAMP Receptor Protein (CRP) in the regulation of the lac operon, we successfully constructed a targeted point mutation within the CRP-binding site of the *lacI* promoter region on the chromosome of the *E. coli K-12* strain BW25113. The genetic modification was performed using a Counter-Selection *E. coli* Chromosome Modification Kit, which enables precise, scarless genome editing. The objective was to disrupt the native CRP-binding site to create a mutant strain designated as BW25113 *lacI*\*.

Following the mutagenesis procedure, several individual colonies were selected and screened. Genomic DNA was isolated from candidate clones (**Figure 4A**), and the targeted *lacI* promoter region was amplified by PCR and subjected to DNA sequencing (**Figure 4B**). The sequencing chromatogram (**Figure 4C**) conclusively confirmed the introduction of the designed nucleotide substitution(s) at the specific sites within the CRP-binding region, thereby validating the successful disruption of the binding site.

**Figure 4.**
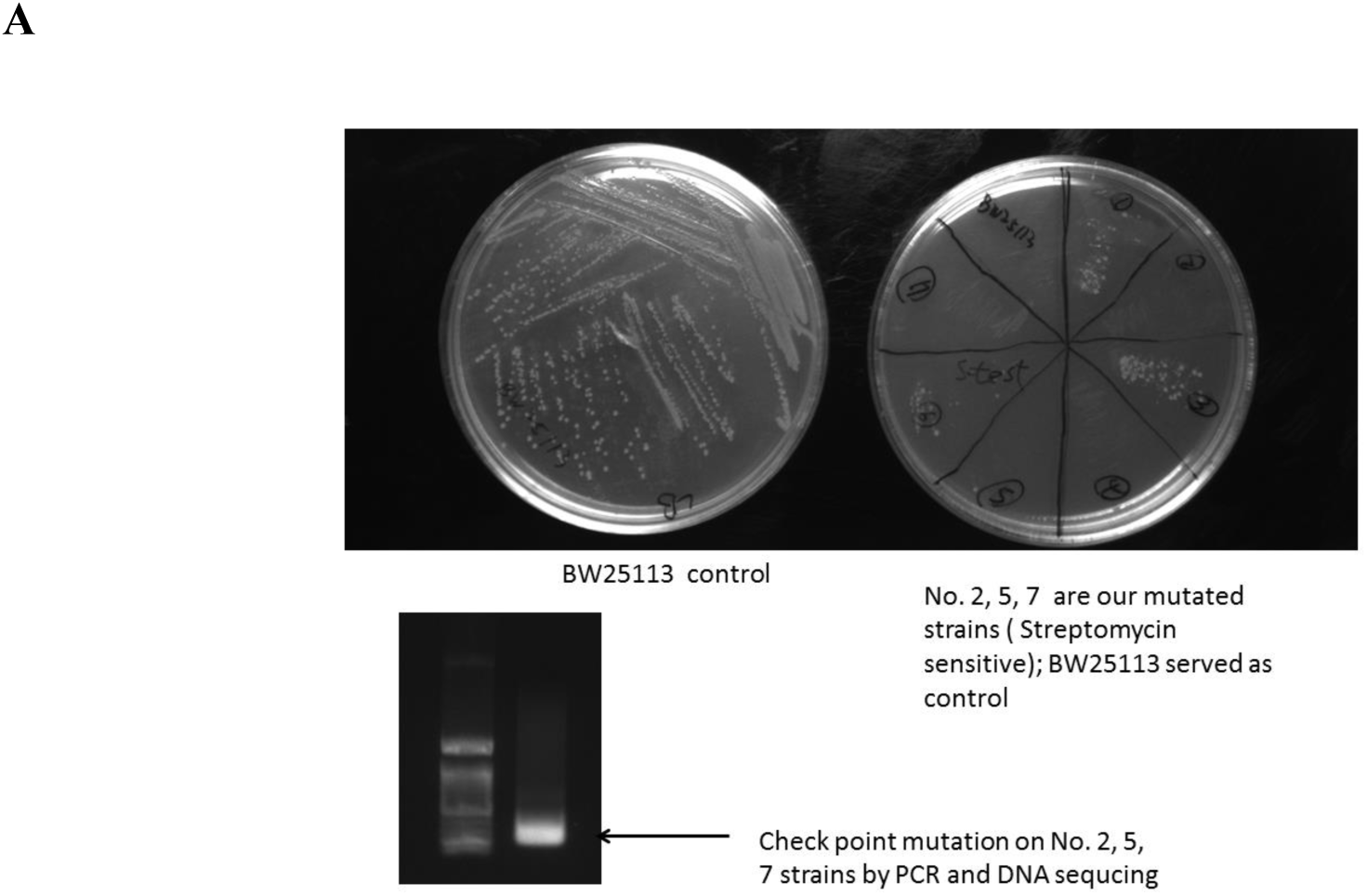
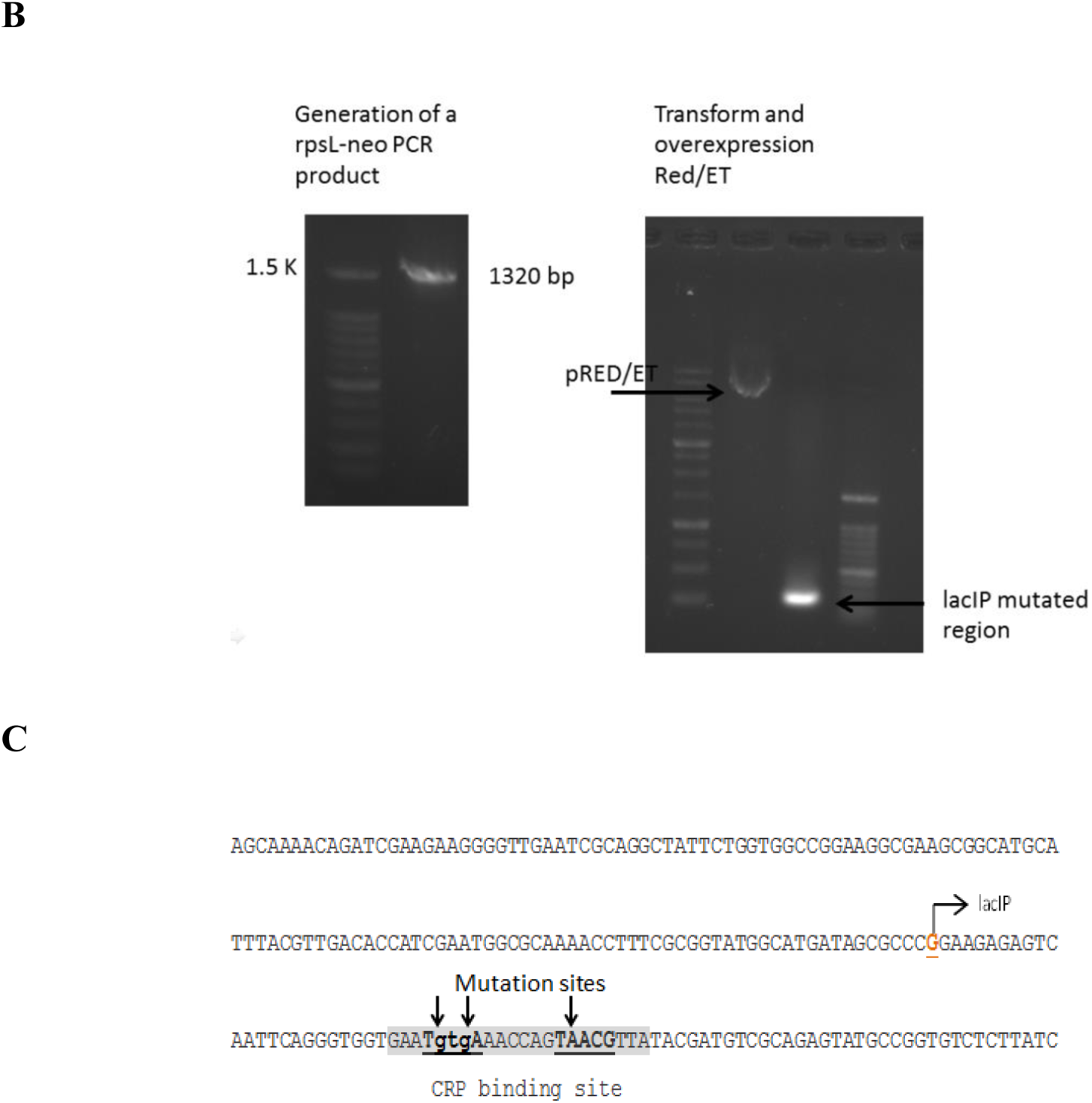
Construction and verification of the lacI promoter CRP-binding site mutant. **A.** Selection of candidate *E. coli* colonies following mutagenesis. **B.** PCR amplification of the targeted *lacI* promoter region from genomic DNA of candidate clones. **C.** DNA sequencing chromatogram of the mutated region, confirming the specific nucleotide substitutions (indicated by the arrow) that disrupt the CRP-binding site.

These results confirm the successful construction of the *E. coli* BW25113 *lacI* mutant strain. This engineered strain provides a valuable tool for further studies aimed at elucidating the CRP-dependent regulatory mechanisms of the *lac* operon, independent of the canonical Lac repressor-mediated regulation.

### cAMP-Dependent Expression of *lacZ* and *lacI* in WT and BW25113 *lacI*\*

To delineate the role of the CRP-binding site in the autoregulation of *lacI* and its subsequent effect on the *lac* operon, we investigated the transcriptional response of both *lacZ* and *lacI* to varying cAMP concentrations in the wild-type (WT) BW25113 and the isogenic *lacI* promoter mutant BW25113 *lacI*\*. Following growth in LB to mid-log phase, cells were transferred to a minimal M9 medium with glycerol as the sole carbon source and induced with a gradient of cAMP for 30 minutes before RNA extraction and analysis. The results revealed a critical disruption in the regulatory circuit in the mutant strain. In the WT, as expected, increasing cAMP concentrations led to a substantial induction of *lacZ* expression (**Figure 5A**). Conversely, in the BW25113 *lacI*\*, this robust induction was markedly attenuated, demonstrating an impaired capacity to fully activate *lacZ* transcription even under high cAMP conditions. The *lacI* gene deletion strain (Δ*lacI*) was used as a control. Notably, quantification of lacI transcript levels revealed that the mutation in its own promoter’s CRP-binding site led to constitutive expression of the lacI repressor, which was unaffected by changes in cAMP levels (**Figure 5B**). This finding confirms that CRP functions as a direct transcriptional repressor of *lacI*.

**Figure 5.**
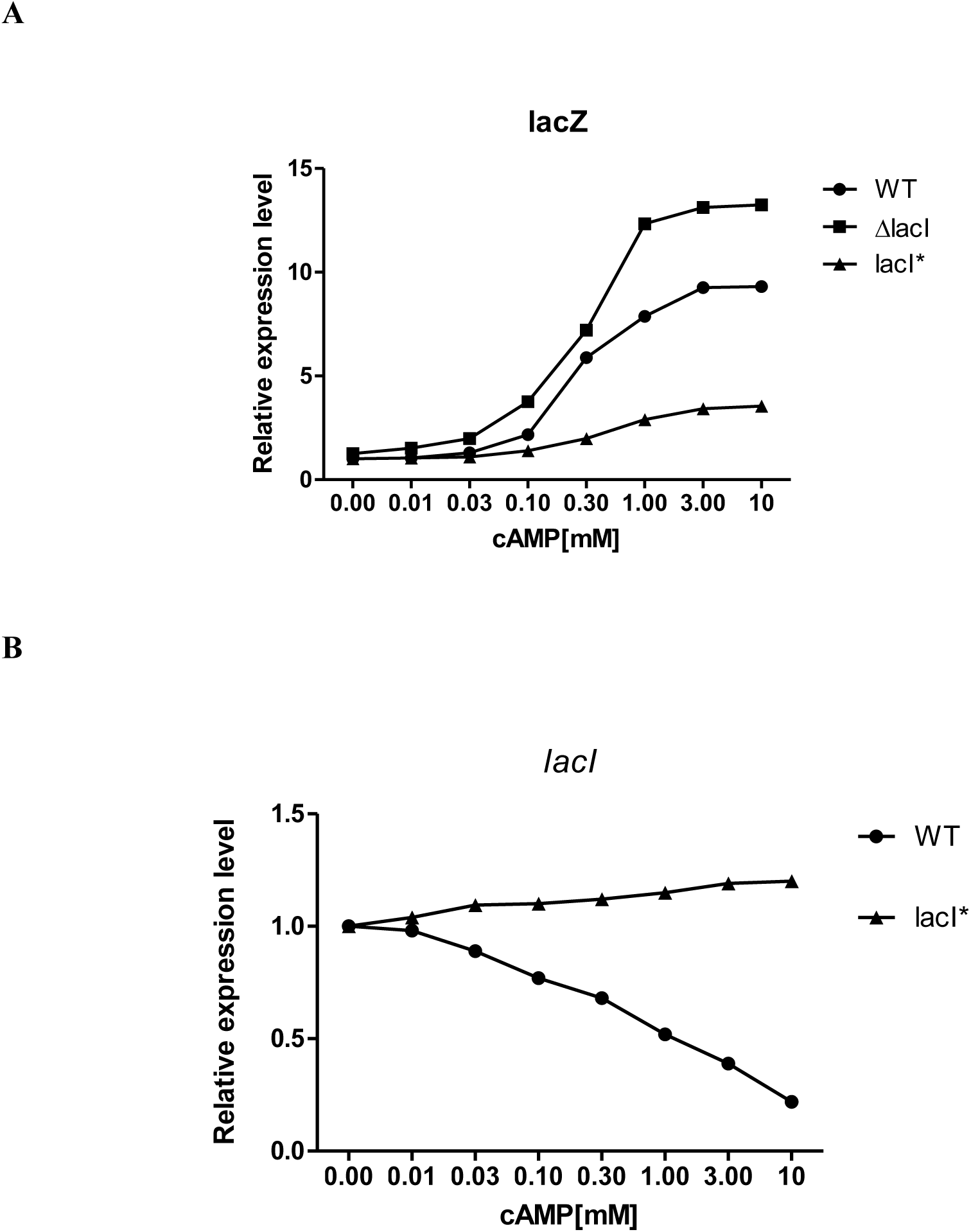
A CRP-mediated feed-forward loop optimizes *lac* operon induction. **A.** *lacZ* and **B**. *lacI* expression in WT, *lacI* promoter mutant *lacI*\*, and Δ*lacI* strains under varying cAMP. The mutant’s constitutive *lacI* expression and attenuated *lacZ* induction reveal a dual role for CRP.

Collectively, these data demonstrate that CRP-mediated repression of *lacI* is essential for the full induction of the *lac* operon. We propose that this regulatory architecture constitutes a Coherent Type 4 Feed-Forward Loop (FFL) involving CRP, LacI, and LacZ. In this model, during growth on dual carbon sources such as glucose and lactose, the depletion of glucose lifts catabolite repression, resulting in increased cAMP and CRP activation. This dual action creates a synergistic effect that potentiates the induction of the *lac* operon. We hypothesize that this FFL enhances the sensitivity for detecting environmental lactose upon glucose exhaustion. In the absence of this circuit, as in our mutant, the constitutive levels of LacI would necessitate a higher concentration of lactose to relieve repression, potentially delaying the metabolic switch to lactose utilization.

### Growth Phenotype of the *lacI* Promoter Mutant in a Glucose-Lactose Diauxic Shift

To assess the physiological impact of disrupting CRP-mediated regulation of the *lacI* promoter, we analyzed the growth of E. coli BW25113 (WT), the isogenic lacI promoter mutant (BW25113 *lacI*\*), and a Δ*lacI* control in minimal medium containing glucose and lactose. As expected, the WT exhibited a characteristic diauxic growth curve, with a distinct pause followed by resumption of growth upon switching from glucose to lactose metabolism. In stark contrast, the BW25113 *lacI*\* mutant displayed a significantly prolonged diauxic lag phase, requiring substantially more time to resume growth and reach a maximum cell density (**Figure 6A)**. This phenotype is consistent with our molecular data: the constitutive expression of the LacI repressor from the mutated promoter, which is unresponsive to cAMP-CRP, maintains repression of the *lacZYA* operon. Consequently, a higher concentration of lactose is required to achieve allosteric relief of this repression, thereby delaying the induction of the lactose utilization genes and extending the lag phase. As predicted, the Δ*lacI* control strain, which lacks the repressor entirely, eliminated the diauxic pause, transitioning directly to lactose metabolism without a detectable lag. Collectively, these growth data directly demonstrate that the CRP-mediated feed-forward loop is not merely a theoretical construct but is critical for ensuring a rapid and sensitive metabolic transition, optimizing fitness in environments with shifting carbon sources.

**Figure 6.**
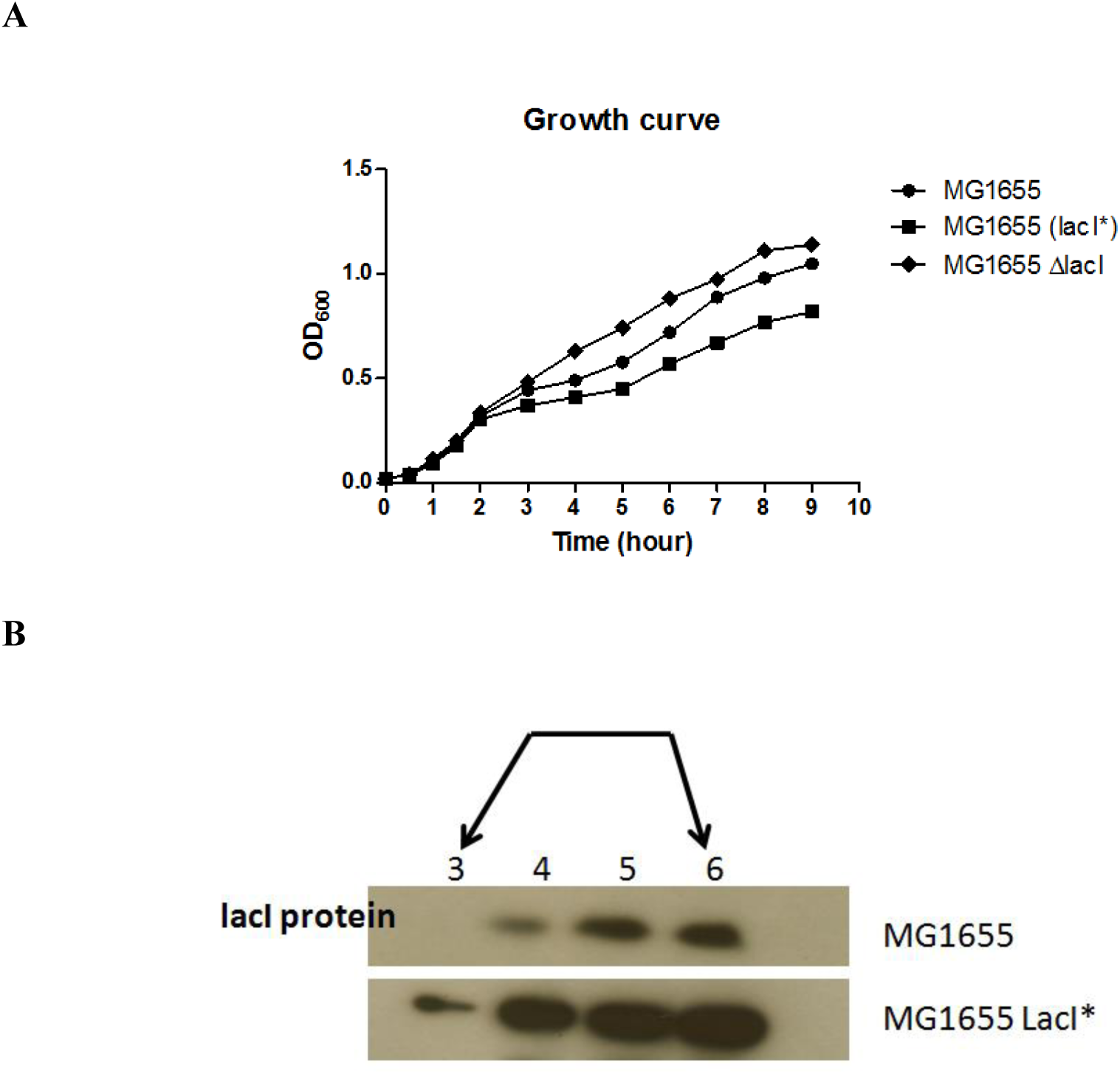
Elevated LacI repressor levels underlie the prolonged diauxie in the lacI promoter mutant. **A.** Growth curves in glucose-lactose medium show a significantly extended diauxic lag in the BW25113 *lacI\** mutant compared to WT and Δ*lacI* strains. **B.** Corresponding LacI protein immunoblot at selected timepoints (hours 3-6) reveals constitutive high-level accumulation of the repressor in the mutant, which maintains repression of the lac operon and delays lactose utilization.

Protein expression analysis was performed on samples collected at time points 3, 4, 5, and 6 hours in Figure 6A during the glucose-lactose diauxic growth (**Figure 6B)**. This analysis confirmed that in the BW25113 *lacI*\* mutant, where *lacI* expression escapes CRP-mediated repression, the cellular concentration of the LacI repressor protein was significantly higher than in the wild-type BW25113 strain. We posit that this accumulated LacI pool consequently imposes a stronger repression on the *lacZYA* operon, ultimately delaying the transition to lactose metabolism, as observed in the extended diauxic lag. To further substantiate this, we will quantify β-galactosidase activity to directly measure the functional outcome on *lacZ* expression. Furthermore, we hypothesize that the CRP-mediated repression of lacI in the wild-type strain serves a critical sensitizing function. By maintaining lower basal levels of LacI, the wild-type BW25113 is predicted to initiate growth on lactose at a significantly lower external lactose concentration compared to the BW25113 *lacI*\* mutant. A lower concentration of the inducer (lactose) is required to relieve the weaker repression in the wild-type, enabling a more sensitive and rapid genetic response.

## Discussion and Conclusion

This study demonstrated that E. coli can establish a control system for carbon source utilization in an effectively enriched environment with double-carbon sources (e.g., glucose and lactose) via Co-4 FFL, formed with CRP, LacI, and *lacZ* (**Figure 7)**.

**Figure 7.**
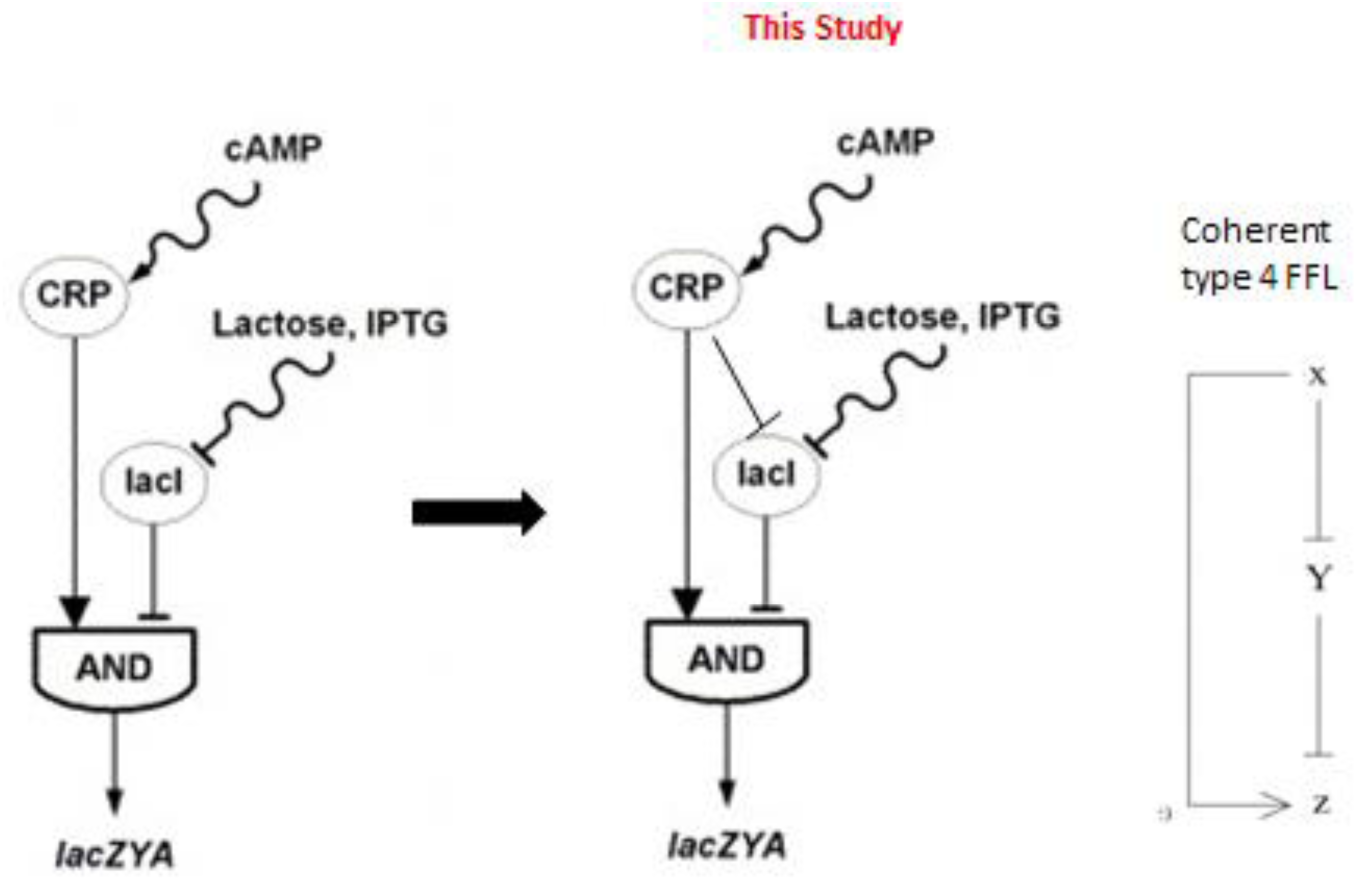
New regulation model of CRP-LacI-*lacZ* Coherent type 4 feed-forward loop in *Escherichia coli*.

In this system, a decrease in glucose concentration can induce the formation of the cAMP-CRP complex, thereby decreasing the concentration of LacI. This system also induces *lacZ* gene expression at an appropriate time and largely increases sensitivity to lactose concentration. Lactose metabolism occurs to produce energy when a small amount of lactose is left and glucose is consumed. On the contrary, if this system is damaged, the Lac operon in *E. coli* is stimulated by a high lactose concentration in a double-carbon source environment; hence, lactose consumption is delayed. This result may prolong the time *it takes for E. coli* to adapt in environments with double-carbon sources. *E. coli* may also suffer from a lack of carbon sources because of the failure to metabolize lactose rapidly, resulting in competition for lactose among other bacteria in the intestinal tract of animals.

We also found that a CRP binding site is present in the region of the *lacI* gene promoter. Based on these findings, we re-investigated the gene regulation of the lac operon described in previous studies. With a re-established relationship among CRP, LacI, and *lacZ* gene regulation, the *la*c operon, regulated by double transcription factors (CRP and LacI), may be altered to form a gene regulatory network topology with a feed-forward loop (FFL). In such a gene regulatory relationship, CRP can be used to positively regulate the lacZ gene and negatively regulate the lacI gene. Moreover, LacI repressor (*lacI*) in the lac operon can inhibit the *lacZ* gene. An interdependent regulatory relationship among these three genes is sufficient to form coherent type 4 FFL (Co4 FFL). However, studies have yet to investigate the functions of this pattern of gene regulatory module in *E. coli.* This study aimed to investigate the functions of Co4 FFL formed with the transcription factor CRP and *the lac operon, which are relevant to* lactose metabolism. Moreover, this study elucidated the mechanism by which a control system for carbon sources can be provided for E. coli in an effectively enriched environment by using Co4 FFL, formed by CRP, LacI, and lacZ, when double-carbon sources (e.g., glucose and lactose) are present in the environment.

It suggests that *E. coli* uses CRP to repress lacI expression, increasing the efficiency of metabolizing lactose for energy synthesis under glucose exhaustion in glucose and lactose diauxic growth conditions. The hypothesis was further verified by site-directed mutation of the CRP binding site on the *lacI* promoter region. The function of a CRP-LacI-*lacZYA* coherent type 4 feed-forward loop in *E. coli* was demonstrated by determining the regulatory role of CRP on LacI repressor and the *la*c Operon. Models of regulation of the *lac* operon in *E. coli* by the CRP activator and LacI repressor reveal the importance of metabolism in determining the behavior of genetic regulatory networks.

With a re-established relationship among CRP, *lacI*, and *lacZ* gene regulation, the *lac* operon, regulated by double transcription factors (CRP and LacI), may be altered to form a gene regulatory network topology with a feed-forward loop (FFL). In such a gene regulatory relationship, CRP positively regulates the *lacZ* gene, whereas it negatively regulates the *lacI* gene. Moreover, LacI repressor (*lacI*) in the lac operon can inhibit the *lacZ* gene. An interdependent regulatory relationship among these three genes is sufficient to form CRP–LacI–*lacZ* coherent type 4 FFL (Co4 FFL), suggesting that *E. coli* uses CRP to repress *lacI* expression to increase efficiency in metabolizing lactose for energy synthesis when glucose is already exhausted in the presented glucose and lactose diauxic growth condition.

## Acknowledgements

We want to thank Dr. Steve Busby of the School of Biosciences at the University of Birmingham for donating the pRW50, pDU9, and pDCRP plasmids.

## Reference

1. Mangan S, Zaslaver A, Alon U: The coherent feedforward loop serves as a sign-sensitive delay element in transcription networks. J Mol Biol 2003, 334(2):197–204.

2. Alon U: Network motifs: theory and experimental approaches. Nature reviews Genetics 2007, 8(6):450–461.

3. Matthews BW: The structure of β-galactosidase. Cr Biol 2005, 328(6):549–556.

4. Kimata K, Takahashi H, Inada T, Postma P, Aiba H: cAMP receptor protein – cAMP plays a crucial role in glucose-lactose diauxie by activating the major glucose transporter gene in Escherichia coli. P Natl Acad Sci USA 1997, 94(24):12914–12919.

5. Traxler MF, Chang DE, Conway T: Guanosine 3 ′, 5 ′ –bispyrophosphate coordinates global gene expression during glucose-lactose diauxie in. P Natl Acad Sci USA 2006, 103(7):2374–2379.

6. Inada T, Kimata K, Aiba HJ: Mechanism responsible for glucose-lactose diauxie in Escherichia coli: Challenge to the cAMP model. Genes Cells 1996, 1(3):293–301.

7. Blaiseau PL, Holmes AM: Diauxic Inhibition: Jacques Monod’s Ignored Work. J Hist Biol 2021, 54(2):175–196.

8. Guerra NP: Enhancing Logistic Modeling for Diauxic Growth and Biphasic Antibacterial Activity Synthesis by Lactic Acid Bacteria in Realkalized Fed-Batch Fermentations. Mathematics-Basel 2025, 13(19).

9. Zhang XR, Chen Y, Zhang XY, Zhu YT, Yang JX, Gong GZ: Comparative transcriptomic analysis reveals a potential link between sugar transporters and the diauxic growth of YT175 on inulin. Int J Biol Macromol 2025, 298.

10. Narang A, Pilyugin SS: Bacterial gene regulation in diauxic and non-diauxic growth. J Theor Biol 2007, 244(2):326–348.

11. Araki N, Niikura Y, Miyauchi K, Kasai D, Masai E, Fukuda M: Glucose-Mediated Transcriptional Repression of PCB/Biphenyl Catabolic Genes in RHA1. J Mol Microb Biotech 2011, 20(1):53–62.

12. Yang CD, Huang HY, Shrestha S, Chen YH, Huang HD, Tseng CP: Large-Scale Functional Analysis of CRP-Mediated Feed-Forward Loops. Int J Mol Sci 2018, 19(8).

13. Manso I, García JL, Galán B: gene expression is regulated by catabolite repression mediated by the cAMP-CRP complex. Microbiol-Sgm 2011, 157:593–600.

14. Baba T, Ara T, Hasegawa M, Takai Y, Okumura Y, Baba M, Datsenko KA, Tomita M, Wanner BL, Mori H: Construction of Escherichia coli K-12 in-frame, single-gene knockout mutants: the Keio collection. Mol Syst Biol 2006, 2:2006 0008.

15. Hollands K, Busby SJ, Lloyd GS: New targets for the cyclic AMP receptor protein in the Escherichia coli K-12 genome. FEMS Microbiol Lett 2007, 274(1):89–94.

16. Lin HH, Hsu CC, Yang CD, Ju YW, Chen YP, Tseng CP: Negative Effect of Glucose on ompA mRNA Stability: a Potential Role of Cyclic AMP in the Repression of hfq in Escherichia coli. J Bacteriol 2011, 193(20):5833–5840.

17. Gaudreault M, Gingras ME, Lessard M, Leclerc S, Guerin SL: Electrophoretic mobility shift assays for the analysis of DNA-protein interactions. Methods Mol Biol 2009, 543:15–35.

18. Chen YG, Lin YCD, Luo YJ, Cai XX, Qiu P, Cui SD, Wang Z, Huang HY, Huang HD: Quantitative model for genome-wide cyclic AMP receptor protein binding site identification and characteristic analysis. Brief Bioinform 2023, 24(3).

19. Chng C, Lum AM, Vroom JA, Kao CM: A key developmental regulator controls the synthesis of the antibiotic erythromycin in Saccharopolyspora erythraea. Proc Natl Acad Sci U S A 2008, 105(32):11346–11351.

20. Karr EA: The methanogen-specific transcription factor MsvR regulates the fpaA-rlp-rub oxidative stress operon adjacent to msvR in Methanothermobacter thermautotrophicus. J Bacteriol 2010, 192(22):5914–5922.

21. Zianni M, Tessanne K, Merighi M, Laguna R, Tabita FR: Identification of the DNA bases of a DNase I footprint by the use of dye primer sequencing on an automated capillary DNA analysis instrument. J Biomol Tech 2006, 17(2):103–113.

22. Chen YG, Mao RB, Xu JT, Huang YX, Xu JY, Cui SD, Zhu ZH, Ji X, Huang SH, Huang YZ et al: A Causal Regulation Modeling Algorithm for Temporal Events with Application to ‘s Aerobic to Anaerobic Transition. Int J Mol Sci 2024, 25(11).

23. Heermann R, Zeppenfeld T, Jung K: Simple generation of site-directed point mutations in the Escherichia coli chromosome using Red(R)/ET(R) Recombination. Microbial cell factories 2008, 7:14.

24. Kolb A, Busby S, Buc H, Garges S, Adhya S: Transcriptional regulation by cAMP and its receptor protein. Annual review of biochemistry 1993, 62:749–795.

25. Lawson CL, Swigon D, Murakami KS, Darst SA, Berman HM, Ebright RH: Catabolite activator protein: DNA binding and transcription activation. Current opinion in structural biology 2004, 14(1):10–20.

26. Blaszczyk U, Polit A, Guz A, Wasylewski Z: Interaction of cAMP receptor protein from Escherichia coli with cAMP and DNA studied by dynamic light scattering and time-resolved fluorescence anisotropy methods. Journal of protein chemistry 2001, 20(8):601–610.

27. Aiba H, Mori K, Tanaka M, Ooi T, Roy A, Danchin A: The complete nucleotide sequence of the adenylate cyclase gene of Escherichia coli. Nucleic acids research 1984, 12(24):9427–9440.

28. Chen YP, Lin HH, Yang CD, Huang SH, Tseng CP: Regulatory role of cAMP receptor protein over Escherichia coli fumarase genes. J Microbiol 2012, 50(3):426–433.

